# Stability by gating plasticity in recurrent neural networks

**DOI:** 10.1101/2020.09.10.291120

**Authors:** Katharina A. Wilmes, Claudia Clopath

## Abstract

With Hebbian learning ‘who fires together wires together’, well-known problems arise. Hebbian plasticity can cause unstable network dynamics and overwrite stored memories. Unstable dynamics can partly be addressed with homeostatic plasticity mechanisms. Unfortunately, the time constants of homeostatic mechanisms required in network models are much shorter than those measured experimentally.

We propose that homeostatic time constants can be slow if plasticity is gated. We investigate how gating plasticity influences network stability and memories in plastic balanced spiking networks of neurons with dendrites. We compare how different factors such as excitability, learning rate, and inhibition lift the requirements for homeostatic time constants. We investigate how dendritic versus perisomatic gating allows for different amounts of weight changes in stable networks. We suggest that the compartmentalisation of pyramidal cells enables dendritic synaptic changes while maintaining stability. We show that spatially restricted plasticity improves stability. Finally, we compare how different gates protect memories.

**Significance statement:** How does the brain maintain stable neural activity in the presence of synaptic changes? This question has been studied extensively in the past, but we argue that one crucial aspect is missing in previous studies. While all theoretical work has assumed plasticity to be *on* all the time, plasticity is in fact heavily gated. In this light, we must reconsider the theories on stability and homeostasis of neural activity. In particular, theoretical studies show that neural networks undergoing plasticity require fast compensatory homeostatic mechanisms to be stable. However, experimentally measured homeostatic processes operate on much slower time scales. We studied how the gating of plasticity can improve network stability and thereby reduce the discrepancy in the homeostatic time constant between models and experiments.

## 1 Introduction

Hebbian plasticity is considered to be the neural hallmark for learning and memory. It enables the formation of cell assemblies as it strengthens connections between cells with correlated activity. On the downside, correlations between cells are increased even further with Hebbian plasticity. Theoretically, such a positive feedback loop leads to undesired unstable runaway activity (Abbott and Nelson, 2000). Cortical cells, however, fire at low rates in an asynchronous irregular manner. It is therefore unclear how neural activity in the functioning brain remains stable despite Hebbian plasticity. To resolve this dilemma, it has been suggested that homeostatic processes keep the network activity stable (Turrigiano and Nelson, 2004). Homeostatic processes, such as homeostatic scaling (Turrigiano *et al.*, 1998; Turrigiano, 2017; Desai *et al.*, 2002; Goel and Lee, 2007; Keck *et al.*, 2013; Maffei and Turrigiano, 2008) or inhibitory plasticity (Woodin and Maffei, 2010; Goel and Lee, 2007; Kuhlman *et al.*, 2013; Keck *et al.*, 2011; Chen *et al.*, 2011; Vogels *et al.*, 2011; van Versendaal and Levelt, 2016; Li *et al.*, 2014), counteract increases in the network activity, but it has been proposed that they might be insufficient to keep the network activity stable for the following reason: these processes operate on a timescale of hours or days (Keck *et al.*, 2017; kaneko *et al.*, 2008a; Greenhill *et al.*, 2015; Kaneko *et al.*, 2008b; Watt and Desai, 2010), but theoretical models require homeostatic mechanisms that act on the same timescale as Hebbian plasticity or faster (Zenke *et al.*, 2013; Zenke *et al.*, 2017; Toyoizumi *et al.*, 2014; Litwin-Kumar *et al.*, 2016; Miller and MacKay, 1994; van Rossum *et al.*, 2000). Zenke *et al.*, 2017, therefore, proposed that there must be a fast compensatory mechanism. Such a mechanism could modulate plasticity itself (Naumann and Sprekeler, 2020). Models requiring fast mechanisms typically assume that plasticity is continuously happening (e.g. Litwin-Kumar *et al.*, 2016; Zenke *et al.*, 2013). In contrast, in the brain plasticity is highly regulated by different neuromodulators (Couey *et al.*, 2007; Bissière *et al.*, 2003; Lin *et al.*, 2003; Pawlak *et al.*, 2010; Seol *et al.*, 2007; Zhang *et al.*, 2009; Brzosko *et al.*, 2015), astrocytes (Valtcheva and Venance, 2016), and inhibitory interneurons (Artinian and Lacaille, 2018; Kuhlman *et al.*, 2013; Letzkus *et al.*, 2011). These different regulators of plasticity can slow down, speed up, gate, or flip plasticity. They differ in their temporal and spatial precision and hence enable rigorous plasticity control. We, therefore, explore how different regulators or gates affect plasticity and stability. Specifically, we study using computational modelling whether gating of plasticity can lower the requirements for time constants of homeostatic plasticity to values that are more in line with experimentally observed homeostatic processes without strongly impairing plasticity of synaptic connections.

## 2 Results

### 2.1 Balanced spiking neural network with 2-compartment pyramidal cells

To study how different modulators of plasticity affect stability and plasticity, we built a balanced recurrent network of 1000 excitatory pyramidal cells (E) and 250 inhibitory cells (I, Fig. 1). To explore perisomatic and dendritic gating separately, we modelled the pyramidal cells with two compartments, one for the soma and one for the dendrite (Fig. 1a,b, Naud and Sprekeler, 2018). The somatic compartment represents the perisomatic region, i.e. the soma and the proximal basal and apical dendrites, which contains the perisomatic synapses. The dendritic compartment represents the distal apical dendrites, which we will refer to as the dendrite, which contains the dendritic synapses. Both populations receive Poisson spike trains as external inputs. Before implementing plasticity in our model, we made sure that the network is in the asynchronous irregular regime (Fig. 1 c,e,f,g), due to a balance between excitation and inhibition. That is, strong excitatory recurrent inputs were balanced by strong inhibitory feedback (inhibition-stabilized regime, Tsodyks *et al.*, 1997). On the single-cell level, this is reflected in large excitatory and inhibitory currents, which cancel each other on average (Fig. 1d).

**Figure 1:**
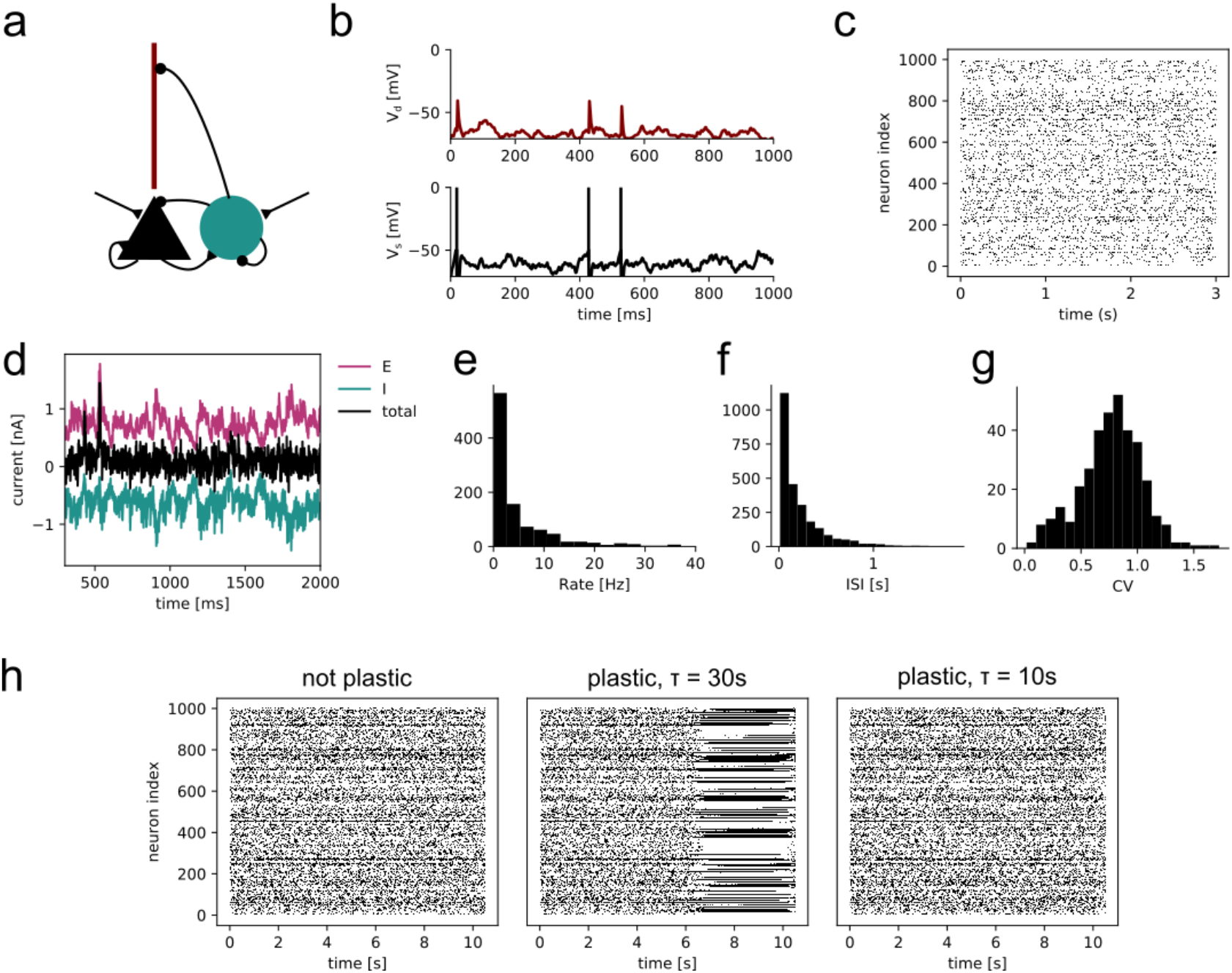
Balanced spiking neural network with 2-compartment pyramidal cells. a: The network consisted of 1000 recurrently connected 2-compartment pyramidal cells (triangle and stick), and 250 recurrently connected inhibitory cells (circle). Both the excitatory and the inhibitory population receive external Poisson inputs (black arrows). b: Somatic (black) and dendritic (red) voltage traces from one example pyramidal cell. c: Raster plot of excitatory cell activity in the network. d: Example currents from one example pyramidal cell. It receives large E (magenta) and I (cyan) currents which cancel on average (black). e: Distribution of excitatory firing rates. f: distribution of excitatory interspike intervals. g: Distribution of coefficient of variation (CV) of the interspike intervals. e-g indicate that the network is in a balanced state. h: Raster plots of excitatory network activity in a network without plasticity (left), with plasticity and a homeostatic time constant *τ* =30ms, and with plasticity and a homeostatic time constant *τ* =10ms.

To test the effect of plasticity in our network, we added a standard triplet STDP rule (Pfister and Gerstner, 2006; Clopath *et al.*, 2010; Bono *et al.*, 2017) to the excitatory connections. As this form of plasticity is Hebbian, it can lead to an explosion of activity in recurrent networks (Keck *et al.*, 2017; Abbott and Nelson, 2000). To keep the activity of the network in the balanced state despite ongoing plasticity, we included homeostatic plasticity (Keck *et al.*, 2017; Abbott and Nelson, 2000). Following previous work (Bienenstock *et al.*, 1982; Pfister and Gerstner, 2006; Clopath *et al.*, 2010; Zenke *et al.*, 2013), the homeostatic process in our network monitored the postsynaptic firing rate and adjusted long-term depression (LTD) to keep the neurons at their target firing rate. The time constant *τ* of this homeostatic process is critical for stability as it determines how quickly the homeostatic process reacts to changes in firing rate. If *τ* is too large, the homeostatic plasticity cannot compensate for the correlation-based weight changes and the network activity explodes (Fig. 1h middle). When *τ* is sufficiently small, the homeostatic plasticity maintains stability (Fig. 1h right). The time constants required for stability in models are orders of magnitude smaller than those of homeostatic processes in the brain. Therefore, we explored in our model whether gating plasticity can allow for larger homeostatic time constants, to reconcile stability constraints with experimental data (Zenke *et al.*, 2017; Turrigiano *et al.*, 1998).

### 2.2 Gating plasticity loosens the requirements for fast homeostatic processes

We next explored how different forms of plasticity modulation affect the required homeostatic time constant in our model. One gate we explored was the excitability of individual neurons to model the fact that neuromodulation can change the size or the duration of excitatory postsynaptic potentials (EPSPs) (Rasmusson, 2000). In our two-compartment leaky-integrate-and-fire neuron model, we modelled excitability as a factor *τ* multiplied to the excitatory synaptic currents (Eq. 1,2, the superscript *d* indicates the dendritic variable). A second gate was *inhibition.* We modelled its modulation by changing the inhibitory conductance ***g_I_*** (Eq. 12). The somatic and dendritic voltage (*V_s_* and *V_d_* respectively) was therefore modelled as

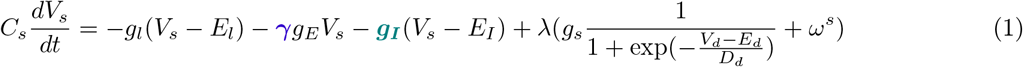

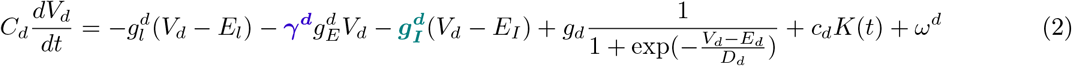

where *C_s_/_d_* is the somatic/dendritic capacitance, *g_I/E_* is the inhibitory/excitatory conductance (index d indicates the dendritic variables), *E_l_/E_I_* is the reversal potential of the leak/the inhibitory synapses (note that the reversal potential for the excitatory synapses is 0 mV and hence omitted), *g_s_* is the coupling from the dendrite to the soma, 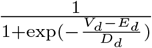 is a nonlinear term for the dendritic calcium dynamics (see Methods), *ω^s/d^* is the somatic/dendritic adaptation variable, *K* the kernel for the back-propagating action potential current with amplitude *c_d_* (Naud and Sprekeler, 2018). λ ensures that the somato-dendritic coupling and adaptation are the same as in the model of Naud and Sprekeler, 2018 (see Methods).

A third gate we explored was the *spiking threshold v_θ_*. When the somatic membrane potential *V_s_* reaches a threshold *v_θ_*, the neuron fires a spike. Spike times *t^f^* are therefore defined as *t^f^*: *V_s_*(*t^f^*) > *v_θ_*. These three gates modulate plasticity indirectly by modulating the activity of the network. A fourth gate was *learning rate,* which modulates the synaptic weight changes directly, and was modelled as a factor ***η*** in the weight update. Formally, the perisomatic synaptic weight and dendritic synaptic weight 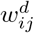 from neuron *j* to neuron *i* changed as:

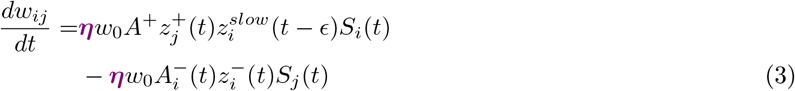

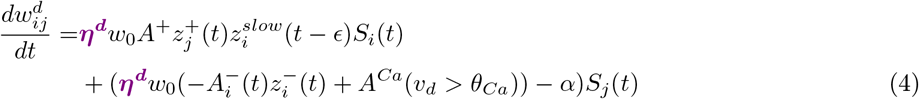

where *ω*_0_ is the initial weight, *A*^+/-^ is the amount of potentiation/depression constants for the triplet rule, *A^Ca^* is the potentiation constant for the Ca^2+^ spike-dependent potentiation, *S_i/j_* is the post-/presynaptic spike train, 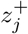 is the presynaptic trace,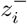 is the postsynaptic trace, 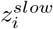 is the postsynaptic trace with slow time constant. *t* – *ϵ* denotes that the value of the trace is taken before the action potential, which happened at time *t. v_d_* is the dendritic membrane potential. *θ_Ca_* is the threshold for Ca^2+^ spike-dependent plasticity (the term *v_d_* > *θ_Ca_* takes values 1 or 0 depending on whether *v_d_* is above the threshold *θ_Ca_*) and *α* is transmitter-induced depression.

To quantify how gating affects stability, we defined the explosion factor as the maximum firing rate in the simulation normalised by the firing rate at the beginning of the simulation, which indicates whether the network is stable (explosion factor close to 1, Fig. 2f) or explodes (explosion factor > 1.5, Fig. 2g). The threshold of 1.5 for a network to be defined as exploding was based on the bimodal distribution of explosion factors (Fig. 2h).

**Figure 2:**
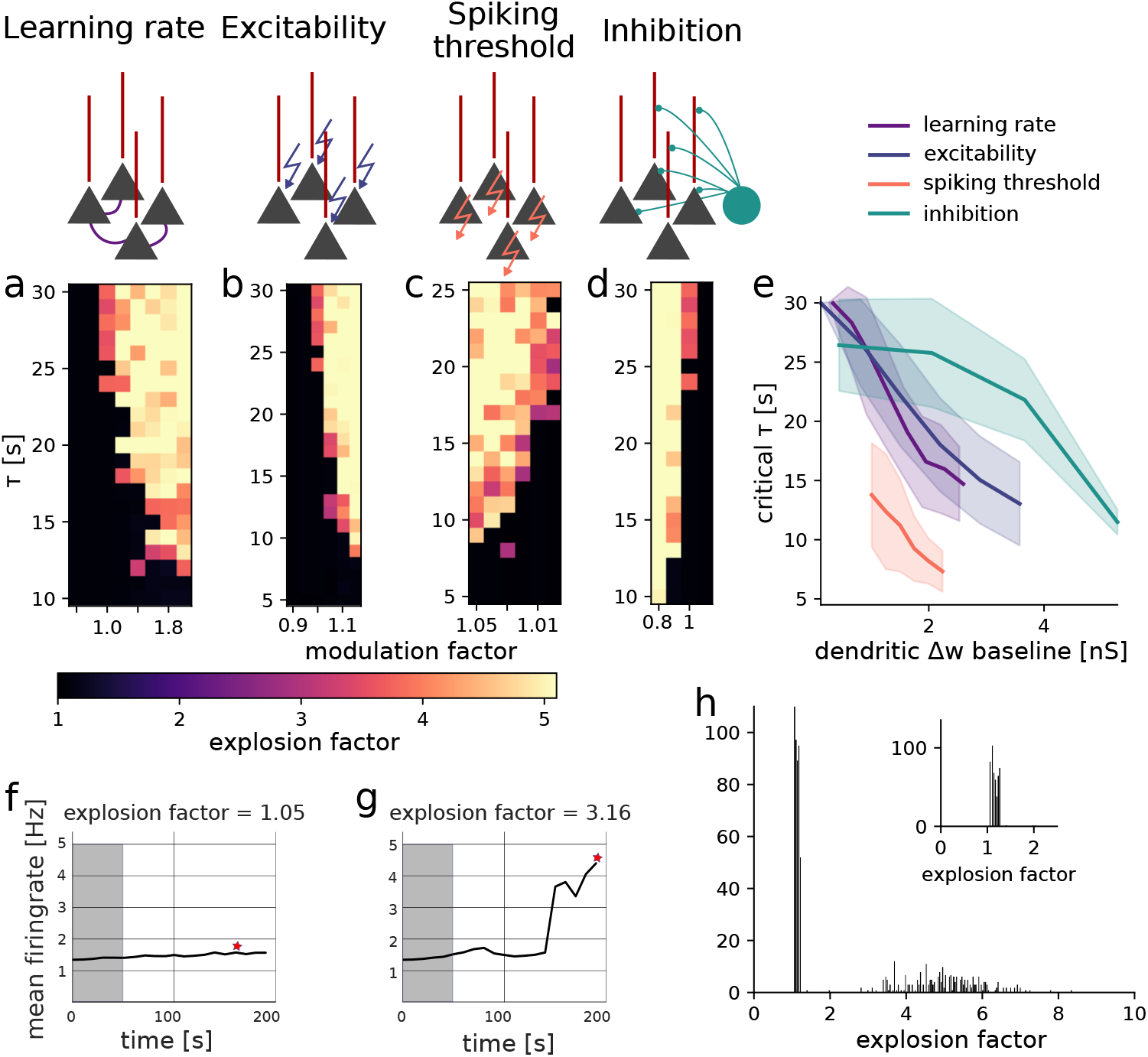
Gating plasticity loosens the requirements for fast homeostatic processes. a-d: Explosion factor as a function of homeostatic time constant *τ* and the respective gate. a: learning rate, b: excitability, c: spiking threshold, d: inhibition. e: comparison of the critical homeostatic time constant *T_crit_* for different gates, plotted as a function of baseline dendritic weight change to allow for comparison. f-g: Illustration of the explosion factor. The star indicates the maximum firing rate of each simulation that was taken for the measurement of the explosion factor. The grey area denotes the reference firing rate at the beginning of the simulation, which was taken to calculate the explosion factor. f: Example network simulation, where the firing rate does not explode (with explosion factor 1.05). g: Example network simulation, where the firing rate explodes (with explosion factor 3.16). h: Distribution of explosion factors. Inset: zoom into the x-axis.

We started by varying learning rate in both the perisomatic and the dendritic compartment (Fig. 2a). Expectedly, we found that with a low learning rate and a large homeostatic time constant *τ*, the network was stable (the black region in Fig. 2a). For higher learning rates, the network activity exploded already at low values of *τ*. This is expected as a higher learning rate increases the rate of synaptic change, which compromises the stability of the network. We defined the largest *τ* at which the network was still stable as the *critical homeostatic time constant τ_crit_* (Fig. 2a). A decrease in learning rate increased this critical time constant *τ_crit_.* Similarly, a decrease in excitability also increased the *τ_crit_* (Fig. 2b). An increase in the spiking threshold has a similar effect as it makes the cells less likely to spike, i.e. less excitable. In these cases, *τ_crit_* decreases, because increasing excitability increases the overall activity in the network, which in turn increases the amount of plasticity. An increase in inhibition on the contrary had the opposite effect on the critical time constant *τ_crit_* (Fig. 2d). Increasing inhibitory inputs decreases firing rates in the network which improves network stability. Therefore, homeostatic mechanisms for network stability can be slower when excitability and learning rate are downregulated or when inhibition is upregulated.

Although these effects were to be expected, qualitatively, our computational model allowed us to compare them quantitatively. We next characterised the different gates by comparing their effects on *τ_crit_.* To compare gates despite their different scales, we defined a common variable. That is, we plotted *τ_crit_* as a function of the total dendritic weight change happening in a stable network (with a *τ* of 5 ms, see Methods, Fig. 2d). This analysis revealed that excitability and learning rate affect the critical time constant *τ_crit_* in a different way than inhibition. *τ_crit_* increases supralinearly as a function of the baseline dendritic weight change for the excitability and learning rate gates, whereas it increases sublinearly for the inhibition gate. The actual dendritic weight change also decreases as a function of the critical homeostatic time constant (Suppl. Fig. S1). To conclude, all gates can improve network stability and lower the requirements for the time constant of homeostatic mechanisms. However, excitability and learning rate have a larger modulating gain than inhibition.

### 2.3 Learning in dendrites facilitates synaptic changes while maintaining network stability

The increase in the critical time constant by gating comes at the cost of the lack of plasticity (measured as the total dendritic weight change, Suppl. Fig. S1). However, pyramidal neurons consist of a soma and a complex ramified structure of dendrites. Interestingly, the majority of excitatory synapses are located on dendrites, electrotonically distant from the soma. Inspired by these observations, we hypothesised that the anatomy of pyramidal cells could enable both plasticity of dendritic synapses and stable somatic activity at the same time. We, therefore, increased the learning rate and the excitability separately for the perisomatic and the dendritic synapses and compared their impact on *τ_crit_.*

We found that increasing plasticity (by increasing learning rate or excitability or decreasing inhibition) in the dendrite compromised the critical time constant *τ_crit_* less than in the perisomatic compartment (Fig. 3). *τ_crit_* was significantly larger for a two-fold increase in the learning rate in the dendrite than for the same increase in the learning rate in the perisomatic compartment (Fig. 3a). Moreover, modulating learning rate only in the dendrite allowed for significantly higher dendritic weight changes at a larger critical time constant (Fig. 3d). Increasing excitability by 15% in the dendrite led to a significantly larger *τ_crit_* than increasing excitability by 15% in the perisomatic compartment (Fig. 3b), while there was no difference in dendritic plasticity between the two conditions (Fig. 3e). Similarly, a 30% decrease in dendritic inhibition maintained a significantly larger *τ_crit_* than the same decrease in perisomatic inhibition (Fig. 3c), while there was no difference in dendritic plasticity (Fig. 3f). Note that we chose a two-fold increase in the learning rate, a 15% increase in excitability, and a 30% decrease in inhibition as these changes lower *τ_crit_* by more than 50% (maximum explored values in Fig. 2e).

**Figure 3:**
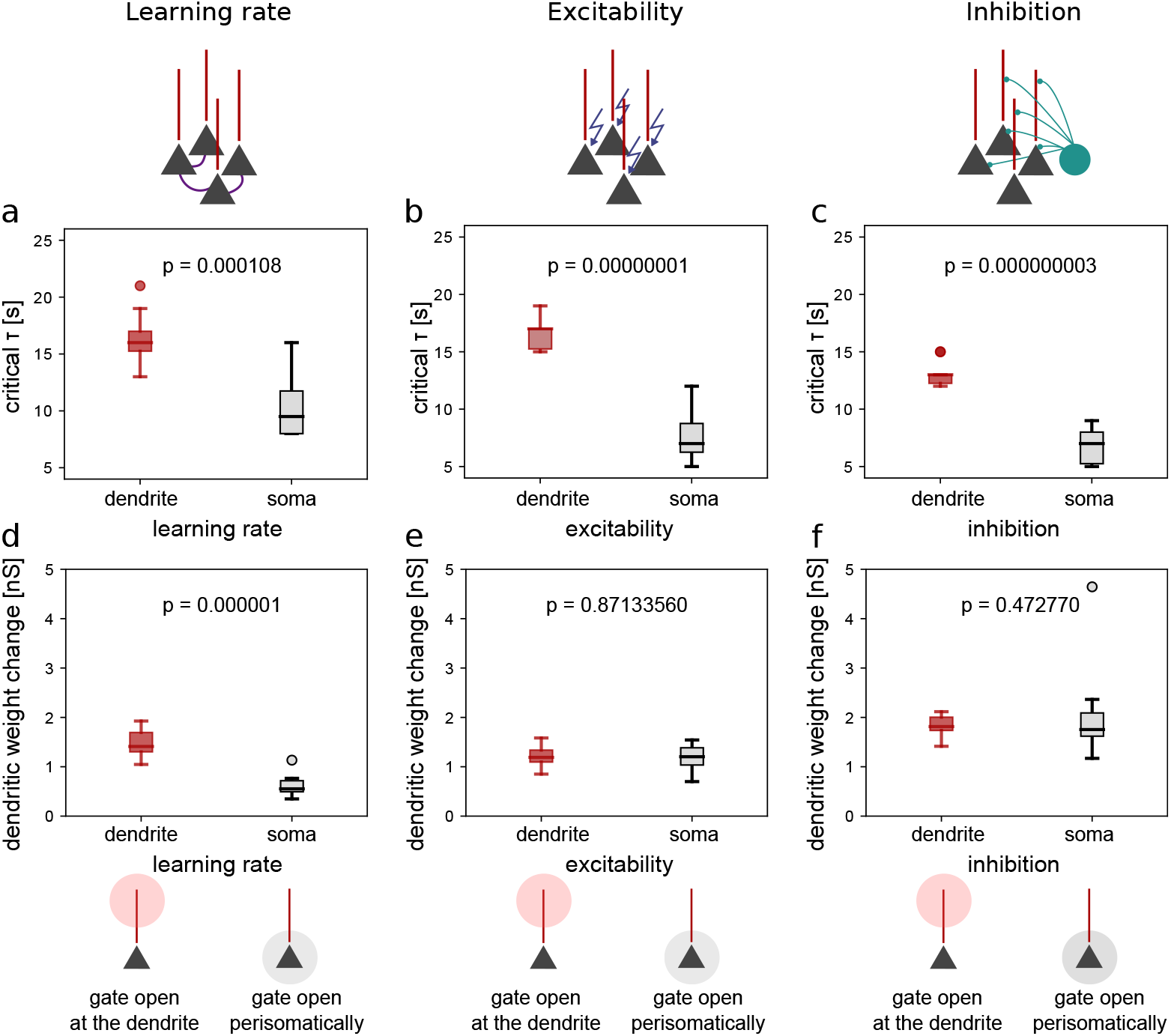
Learning in dendrites helps mitigate the plasticity-stability dilemma. a-c: Distribution of critical homeostatic time constants for gating in the dendritic (red) and in the perisomatic (black) synapses for (a) a two-fold increase in the learning rate, (b) a 15% increase in excitability and (c) a 30% decrease in inhibition. d-f: Distribution of dendritic weight changes for gating in the dendritic (red) and in the perisomatic (black) synapses for (d) a two-fold increase in the learning rate, (e) a 15% increase in excitability and (f) a 30% decrease in inhibition. The rectangles represent the interquartile range (IQR) between first and third quartiles. The thick horizontal lines represent the medians. The whiskers indicate the lowest and highest values within 1.5xIQR from the first and third quartiles, respectively. The circles denote outliers. All p-values were obtained by using the two-sample student’s t-test.

We hypothesised that gating is more effective in the dendrite because of the electrotonic separation of soma and dendrite. To test this hypothesis, we increased the coupling between soma and dendrite. We found that an increase in coupling reduced the critical homeostatic time constant (Suppl. Fig. S2a), in line with our hypothesis. Another property that is special about the dendrite is the dendritic nonlinearity, which induces potentiation of dendritic synapses. Removing the nonlinearity from the model reduced dendritic synaptic weight changes and hence increased the stability of the network (Suppl. Fig. S2b). The combination of synaptic potentiation in the presence of a dendritic nonlinearity and the separation of soma and dendrite hence enables both dendritic weight changes and stable network dynamics.

To further illustrate the effect of a dendritic compartment, we show that critical homeostatic time constants strongly decrease when opening the perisomatic gates in a network of single-compartment neurons (consisting of only a perisomatic compartment, Suppl. Fig. S7).

In summary, by opening the gates for plasticity exclusively in the dendrite, the network can afford slower homeostatic mechanisms, higher network stability, while allowing the same or a higher amount of plasticity as when the gate is open at the perisomatic region.

### 2.4 Spatially precise gating of plasticity enables learning while keeping network activity stable

Neuromodulators were typically thought of as global and diffuse (Schwarz *et al.*, 2015). However, neuromodulatory projections could in principle precisely target specific cell types and subpopulations, depending on their projective field and the receptor channels expressed in their targets. Specific neuromodulation (Totah *et al.*, 2018) could enable plasticity locally when learning requires synaptic adjustments only in a subset of neurons. To test how local gating of plasticity affects the critical time constant *τ_crit_*, we opened the gate for plasticity in only a subpopulation (one-fourth of the neurons) in the network and compared it to opening plasticity in the entire network.

We found that spatially confined plasticity had a much lower impact on the critical time constant than global plasticity. Here, we varied the gates in both the perisomatic and the dendritic compartment. An increase in the learning rate, an increase in excitability, or a decrease of inhibition lowered the critical time constant substantially (Fig. 4 a-c black). Opening these gates in only one-fourth of the neurons lowered the time constant significantly less than opening the gates in the entire network (Fig. 4 a-c green). Therefore, spatially confined gating of plasticity has advantages for network stability beyond enabling precise control.

**Figure 4:**
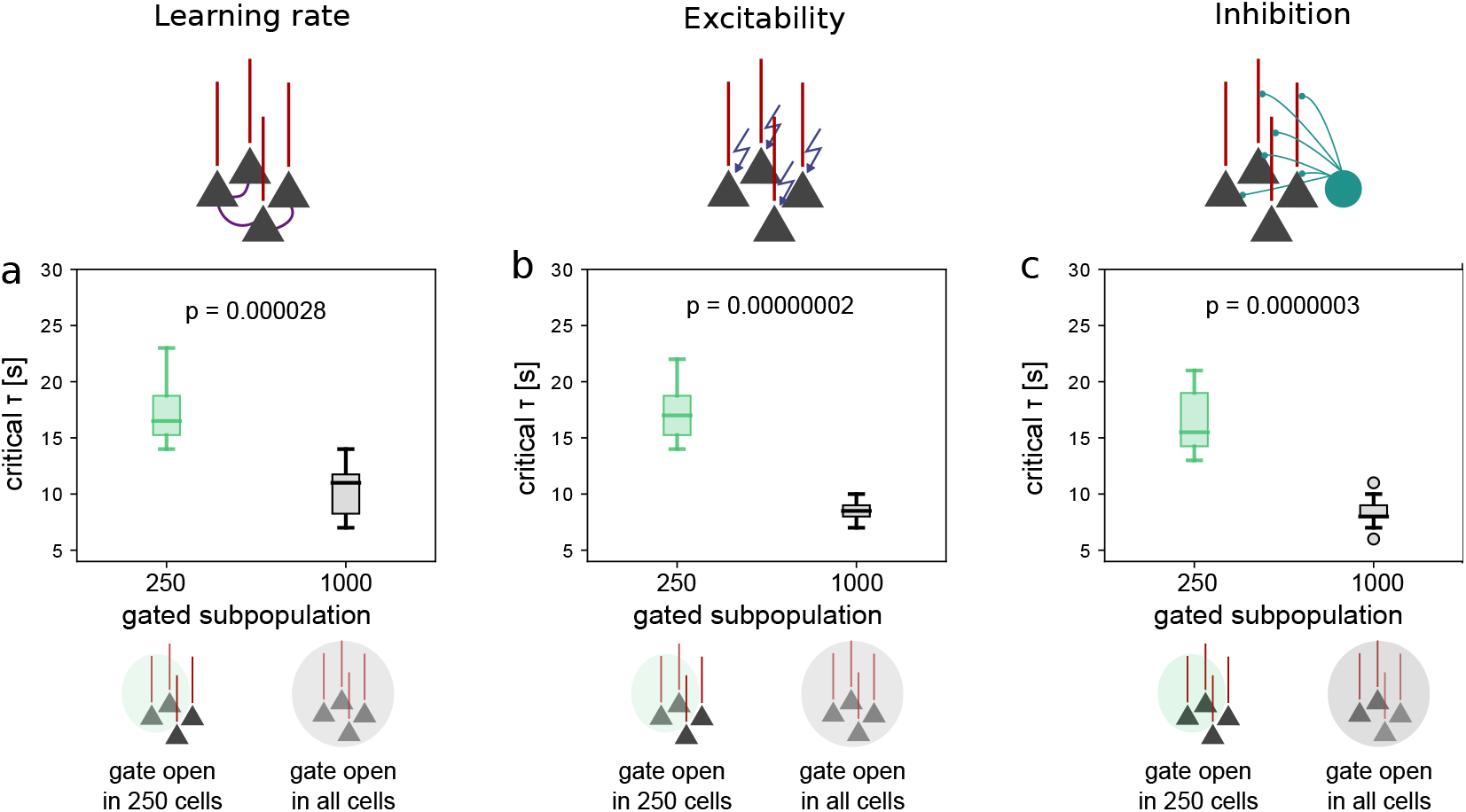
Spatially precise gating of plasticity enables learning while keeping network activity stable. a: Distribution of critical homeostatic time constants for a two-fold increase in the learning rate in a subpopulation of excitatory cells (green) and in the entire network (black). b: Distribution of critical homeostatic time constants for a 15% increase in excitability in a subpopulation of excitatory cells (green) and in the entire network (black). c: Distribution of critical homeostatic time constants for a 20% decrease in inhibition in a subpopulation of excitatory cells (green) and in the entire network (black). The rectangles represent the interquartile range (IQR) between first and third quartiles. The thick horizontal lines represent the medians. The whiskers indicate the lowest and highest values within 1.5xIQR from the first and third quartiles, respectively. The circles denote outliers. p-values were obtained by using the two-sample student’s t-test.

### 2.5 Protection of memories by gating plasticity

In addition to the impact on the stability of network activity, plasticity interferes with the stability of memories. In a plastic network, neural activity patterns lead to synaptic changes, which could overwrite memories that were previously stored in the synaptic connections. Especially when resources are limited, forming new memories can come at the expense of old ones. By gating plasticity, memories can in principle be protected. For example, trivially, by switching off plasticity, old memories are protected. Interestingly, the gates we considered here are not simple all-or-nothing gates but can be continuously modulated. We, therefore, investigated how their modulation affects the maintenance and encoding of memories. Note that we do not use spatially localised gating here.

Before gating plasticity, we first tested how a memory interferes with a previously stored memory in a plastic network. We added inhibitory plasticity (Vogels *et al.*, 2011) to ensure network stability during memory formation and heterosynaptic depression to introduce synaptic competition (see Methods). First, we showed pattern P1 to the network (Fig. 5a). That is, a subpopulation of 100 excitatory neurons received excitatory Poisson input (100 Poisson spike trains with a firing rate of 20 Hz). Afterwards, neurons activated by P1 formed a neural ensemble E1 by increasing their connectivity (Fig. 5a top left). Then, we showed a pattern P2 to the network. P2 is similar to P1 and hence activated a group of neurons E2 that overlapped with the previously formed ensemble. Neurons activated by P2 increased their connectivity. Because the patterns overlapped, synapses were increased at the expense of connections from the old memory (Fig. 5a bottom). Therefore, the new memory interfered with the old memory (Fig. 5 top right). We defined the difference between the mean connection strength of the P1 neurons after memory formation of P1 and the mean connection strength of the P1 neurons at the end of the simulation - after P2 has been learned - as the breakdown of the memory.

**Figure 5:**
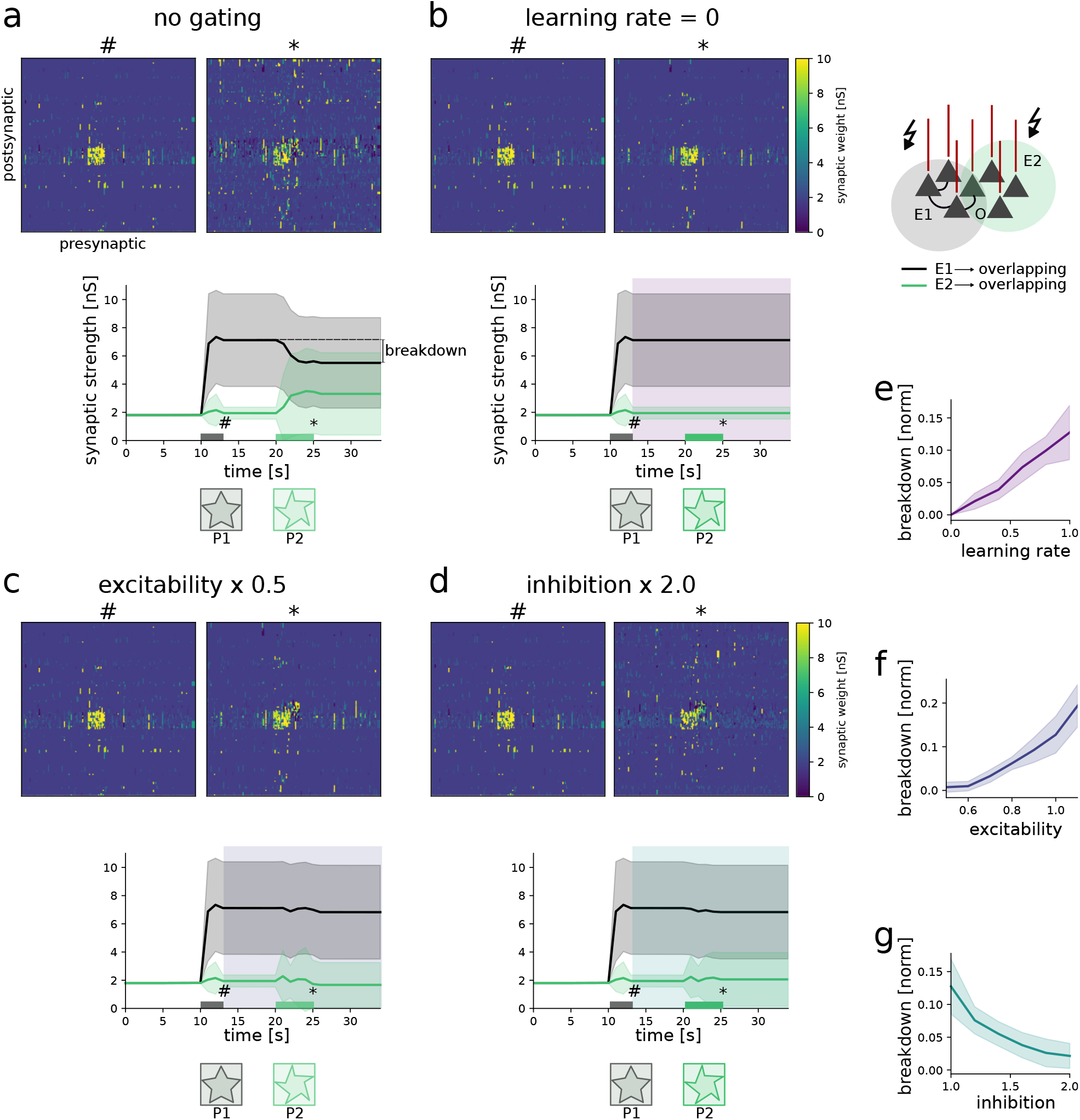
Protection of memories by gating plasticity. a-d: After 10 seconds, we show pattern 1 (P1, grey) to the network for 3 seconds. After a gap of 7 seconds, we show pattern 2 (P2, green) to the network for 5 seconds. Top panels: excitatory weight matrix at two time points of the simulation. Bottom panel: mean synaptic weight from ensemble 1 (E1) to the overlapping region (grey) and from ensemble 2 (E2) to the overlapping region over time as an indication of the strength of the memory of P1 and P2, respectively. a: no gating. b: after P1 is learned, the learning rate is set to 0 (denoted by purple background). c: after P1 is learned, excitability is reduced by 50% (denoted by blue background). d: after P1 is learned, inhibition is doubled (denoted by green background). e-g: breakdown of the memory (see a) as a function of learning rate (e), excitability (f), and inhibition (g).

To test the effect of gating on the protection of memories, we applied the different gates after the first memory is formed. Reducing learning rate to 0 after the first memory is formed trivially protects the memory from being overwritten by the second pattern (Fig. 5b), as this blocked further weight changes. It is less clear how a change in excitability or inhibition affect the storage of the memory. Unlike the learning rate, we cannot modulate excitability or inhibition to their extremes without silencing neural activity. On the one hand, because they decrease neural firing rates, these gates could protect memories by reducing weight changes. On the other hand, by decreasing firing rates, they could also increase LTD as experimentally, low firing rates promote more LTD than LTP (Dudek and Bear, 1992; Sjöström *et al.*, 2001).

We found that a reduction in excitability could indeed protect the memory (Fig. 5c) without permanently silencing the network (Suppl. Fig. S3c). With lower excitability, the stimulated neural ensemble, E2, fired at a lower rate (compare E2 in Suppl. Figs. S3g and S3e), and weights within E2, including those projecting to the overlapping ensemble, O, potentiated less (compare Fig. 5c bottom with Fig. 5a bottom). The new ensemble, E2, hence competed to a lesser extent with the old memory, leading to less memory decay due to heterosynaptic depression. Note that protecting the old memory hence comes at the cost of storing an equally sized representation of P2.

Similar to reduced excitability, increased inhibition could also protect the memory (Fig. 5c) as it reduced firing rates in the network (Suppl. Fig. S3d,h). Notably, the inhibitory plasticity in the network additionally protected the previously formed memory, as it led to increased inhibition of the old memory ensemble E1 (Suppl. Fig. S3i). This further reduced potentiation of synapses from the new ensemble, E2, to the overlapping ensemble, O. We repeated the simulations with an asymmetric inhibitory plasticity rule to show that the results do not depend on the shape of the inhibitory plasticity window (Suppl. Fig. S4).

Because decreased excitability and increased inhibition lower the firing rate of the network, we asked whether this low firing rate induces LTD and hence counteracts the protection of the old memory ensemble. We found that there was no increased LTD within the old memory ensemble due to low firing rates. First, the non-overlapping population of the old ensemble fired at a very low rate (E1-O in Suppl. Fig. S3g,h). There was hence little depression from the non-overlapping population to the overlapping one, as depression happens upon presynaptic spiking. Second, the memory breakdown was weaker at lower excitability and higher inhibition, i.e. at lower firing rates (Fig. 5f,g). The old memory was hence mostly at risk due to heterosynaptic depression. In line with this, the memory breakdown correlated with the maximum mean strength of synaptic connections to the overlapping ensemble O during pattern P2 (Suppl. Fig. S3j-l).

The specificity of the protective effect depends on which gating mechanism is used. For the learning rate, the effect is specific at the synaptic level. For excitability and inhibition, the effect is specific at the cell level. Excitability, inhibition, and learning rate can be modulated at the compartmental level and introduce further specificity there.

When we continuously modulated the gates, we found that the breakdown of the memory increased nonlinearly with both increasing excitability (Fig. 5f) and decreasing inhibition (Fig. 5g), and increased linearly with learning rate (Fig. 5e).

To conclude, all gates can protect memories. Learning rate can be modulated independent of network activity and hence act as a switch for plasticity. Although excitability and inhibition do not modulate plasticity separately, they can both protect memories by reducing activity and weight changes without silencing network activity.

## 3 Discussion

We explored the effect of different gating mechanisms on the stability of neural networks. Using a balanced spiking neural network with 2-compartment pyramidal cells, we showed how gating of plasticity loosens the requirements for fast homeostatic processes. We found that excitability, learning rate, and inhibition affect the critical time constant in different ways. Interestingly, the network was more tolerant towards weight changes, when plasticity gates were opened in the dendrite versus the perisomatic region. Plasticity in dendrites thereby could facilitate learning without compromising the stability of the network.. We also showed that spatially precise gating of plasticity lifts the critical time constant and thereby could locally enable learning while keeping network activity stable. Finally, gating plasticity can protect memories from being overwritten in addition to keeping the network activity stable.

### Examples for the different modulators

Plasticity is highly modulated and gated (Bear and Singer, 1986; Pedrosa and Clopath, 2017; Couey *et al.*, 2007; Bissière *et al.*, 2003; Lin *et al.*, 2003; Pawlak *et al.*, 2010; Seol *et al.*, 2007; Zhang *et al.*, 2009; Brzosko *et al.*, 2015). In this paper, we explored different such modulations of plasticity. First, inhibitory cell types, which target perisomatic and apical dendrites of excitatory cells, can modulate plasticity. It has been shown that disinhibition - the inhibition of inhibitory cells - promotes learning (Letzkus *et al.*, 2011; Kuhlman *et al.*, 2013; Clopath *et al.*, 2016). Dendritic inhibition can influence plasticity directly by affecting depolarizing events in the dendrite, such as back-propagating action potentials and calcium spikes (Larkum *et al.*, 1999; Wilmes *et al.*, 2016). Perisomatic inhibition can modulate plasticity indirectly by decreasing the firing rate of the neuron, as synaptic weight changes depend on neural activity. The modulation of plasticity via inhibition can be both 1) fast, because interneurons can be switched on and off quickly, and 2) local, because they can be precisely targeted by cholinergic (Woody and Gruen, 1987; Metherate *et al.*, 1992; Xiang *et al.*, 1998; Froemke *et al.*, 2007; Froemke *et al.*, 2013) and noradrenergic neuromodulation (Kuo and Trussell, 2011; Martins and Froemke, 2015).

In addition, neuromodulators influence plasticity by regulating neural excitability (acetylcholine and noradrenaline (Xiang *et al.*, 1998; Rasmusson, 2000; Joshi *et al.*, 2016)). For example, acetylcholine binds to muscarinic receptors, which activate a cascade that leads to a decreased permeability of potassium channels (Rasmusson, 2000). This prolongs the duration of EPSPs and thereby increases excitability.

A similar form of neuromodulation is achieved by presynaptic inhibition. A recent theoretical study showed that presynaptic inhibition can act as a fast modulator of plasticity to stabilize network activity (Naumann and Sprekeler, 2020). They showed that presynaptic inhibition is an attractive control mechanism as it depends on network activity and therefore provides a gain control loop. Similar to excitability in our model, the analysis in Naumann and Sprekeler, 2020 shows a supralinear relationship between presynaptic inhibition strength and the critical homeostatic time constant (Fig. 2).

Finally, because many forms of plasticity are NMDAR-dependent (Dudek and Bear, 1992), a modulation of NMDA channels could affect plasticity directly. NMDA channel permeability can be modulated by D-serine, the origin of which is debated (Wolosker *et al.*, 2016), although it was initially thought to be synthesised by astrocytes (Wolosker *et al.*, 2016; Henneberger *et al.*, 2010). Such a direct modulation of plasticity would correspond to modulation of learning rate in our model. A different form of learning rate modulation could be achieved by dendritic inhibition, which is precisely timed to not affect the integration of EPSPs from the dendrite to the soma (Wilmes *et al.*, 2016).

Localised gating could also be achieved by the interplay of multiple mechanisms or network effects. For example, non-specific gating together with the specific feedforward input could lead to specific activity-dependent gating (by a coincidence mechanism).

Hence, plasticity is highly gated and modulated. Depending on the form of modulation, the effect on plasticity can be precisely timed and spatially confined.

### Predictions from the model

We showed that larger synaptic changes are tolerated in dendrites than in the perisomatic region for the same critical time constant (Fig. 3). Therefore, our model predicts that more weight changes should be seen in dendrites. The weaker the dendrite and the soma are coupled, the larger becomes the advantage of the separate dendritic compartment. Hence, we predict that neurons with electrotonically more separate dendrites can undergo more dendritic plasticity.

If plasticity is gated in space and time, i.e. synaptic changes are only locally permitted in limited periods of time, then we would observe that the total amount of synaptic change is not constant, but varies in time and space. The amount of synaptic change averaged over longer periods of time may be constant. When taking averages over shorter periods, we predict that the amount of synaptic change varies significantly over time.

Our model shows that the gates differ in their impact on the critical homeostatic time constant (Fig. 2e). We found that, for inhibition, the critical time constant decreases sublinearly as a function of the resulting increased dendritic weight change. For excitability and learning rate, however, the critical time constant decreases supralinearly. Our model, therefore, predicts that gating plasticity with inhibition allows for a larger critical time constant than gating plasticity with excitability or learning rate. We predict that when inhibition and excitability are separately modulated in an experiment, that the network will lose stability earlier with a change in excitability than with a change in inhibition.

We found that the gates also differ in their ability to protect memories (Fig. 5e-g). Learning rate is the only gate which can completely switch off plasticity to protect the memory. The memory breakdown increased supralinearly with a change in inhibition or excitability, whereas it increased linearly with a change in learning rate. Our model hence predicts that memories break down earlier when inhibition or excitability are modulated than with modulation of learning rate.

### Homeostatic mechanisms and fast compensatory processes

The homeostatic mechanisms which cause the dilemma reported by Zenke *et al.*, 2017 and our paper act on long time scales (hours to days) on the synaptic strengths, as e.g. the BCM sliding threshold and synaptic scaling. They ensure that synaptic weights do not grow unlimited. They can be considered homeostatic because they achieve a certain set point that is stable on average over long time scales. They are feedback controllers, which sense a recent average of the firing rate and adjust weights accordingly. To stabilise Hebbian plasticity, homeostatic mechanisms typically need to be as fast as or faster than the destabilising Hebbian plasticity (Zenke *et al.*, 2017). Therefore, as Zenke *et al.*, 2017 point out, there must be other fast compensatory mechanisms in addition to those slow homeostatic mechanisms. Modelling studies used e.g. inhibitory plasticity with a fast timescale, or heterosynaptic or transmitter-induced plasticity to keep the models stable (Zenke *et al.*, 2015; Litwin-Kumar *et al.*, 2016). Inhibitory plasticity may have a stabilising role (Sprekeler, 2017), but the time scale of inhibitory plasticity appears to be rather slow in comparison to excitatory plasticity (Froemke *et al.*, 2007). Presynaptic inhibition (Naumann and Sprekeler, 2020) or intrinsic plasticity processes that act on the order of minutes (van Welie *et al.*, 2004; Misonou *et al.*, 2004) are good candidates for fast compensatory mechanisms. For any such mechanism, it is however important that it does not destroy the signal or prevent plasticity altogether. To achieve both stability and plasticity, it is important that weight changes can occur. The homeostatic set point of weights should be achieved on average over longer time scales, while allowing temporal deviations from the setpoint (Zenke *et al.*, 2017). The gates, we study here, especially excitability, spike threshold and inhibition could be the target of fast compensatory mechanisms. The point of our study, however, is that plasticity is not always switched on, which is often disregarded in modelling studies. If opening the gates for plasticity is required for synaptic change, networks become much more stable.

### Storing overlapping memories

We showed that all gates protect memories from being overwritten by overlapping neural activity patterns. If the overlapping activity pattern is a new experience that is deemed important enough to be stored in memory, such that the gate for plasticity is opened, the problem of overwriting reappears. Then, further mechanisms are needed to decide whether a memory should be updated to incorporate new features, or whether the new experience is different enough from the memory to be considered a different memory (pattern separation). Sensory discrimination tasks, which reward the successful discrimination of stimuli, show that cells change their selectivity such that the task-relevant stimuli are better represented (Khan *et al.*, 2018; Goltstein *et al.*, 2013; Poort *et al.*, 2015). Therefore, the available pool of neurons or the selectivity of the neurons could change to allow for an equally-sized representation of both stimuli.

### Limitations

Our model provides a comparison between different gating mechanisms. The precise values for the critical homeostatic time constant depend on parameter choices (Suppl. Fig. S6). We simulated a balanced spiking network undergoing spontaneous activity to allow for the comparison of the different plasticity gates. A network which is externally stimulated may have additional requirements for the homeostatic time constant.

In our model, we used one form of homeostatic plasticity, which adjusts LTD based on the postsynaptic firing rate. There are, however, different forms of homeostatic plasticity such as inhibitory plasticity (Woodin and Maffei, 2010) and synaptic scaling (Turrigiano *et al.*, 1998). Inhibitory plasticity also requires fast homeostatic mechanisms when plasticity is not gated (Zenke *et al.*, 2017; Litwin-Kumar *et al.*, 2016). With synaptic scaling as a homeostatic mechanism in our network (Suppl. Fig. S5), gating plasticity increases stability. We, therefore, expect that the gates studied here will similarly lift the requirements for the time scale of inhibitory plasticity and synaptic scaling. It will be interesting to explore the effects of the inhibitory gate on a homeostatic mechanism, which depends on inhibitory plasticity.

### Conclusion

In summary, our study using balanced spiking neural networks with 2-compartment pyramidal cells shows how different forms of gating plasticity increase the stability of neural networks in the presence of plasticity. Our results suggest an important role for dendrites as they can undergo synaptic plasticity with minor effects on network stability. Our results also imply that gating should be locally restricted, supporting the recent finding that neuromodulation may be more specific than initially thought (Totah *et al.*, 2018). Finally, diverse gating mechanisms can protect memories, even if they affect network activity, such as excitability or inhibition.

## 4 Materials and Methods

### Balanced network

We built a recurrent neural network model with *N_E_* =1000 excitatory (E) and *N_I_* =250 inhibitory (I) cells. Both E and I cells received excitatory inputs from a pool of 1000 Poisson processes with a firing rate of 2 Hz and with a connection probability of p=10%. The E cells receive these inputs onto their perisomatic compartment. All neurons were randomly connected. Excitatory cells receive excitatory and inhibitory synapses on both their perisomatic and their dendritic compartment. The connection probability is 10% for all connections except from excitatory cells to excitatory cell’s perisomatic compartment. The connection probability for those connections is 9% to account for the fact that the cells also receive inputs on their dendrites in the two-compartment model. The connection strength of the synapses is chosen such that the network is balanced (see Table 4).

### 2-compartment pyramidal cell model

For the excitatory population, we used a 2-compartment integrate and fire pyramidal cell model with spike-triggered adaptation, adapted from the model by Naud and Sprekeler, 2018 which was originally fitted to data from layer 5 pyramidal cells. It has two coupled membrane equations, one for the soma (*V_s_*, Eq. 1), one for the dendrite (*V_d_*, Eq. 2), modelled as (for clarity we repeat the equations from the main text):

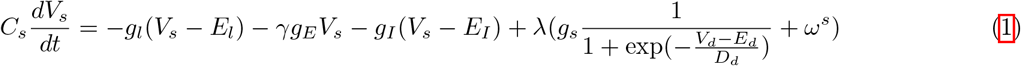

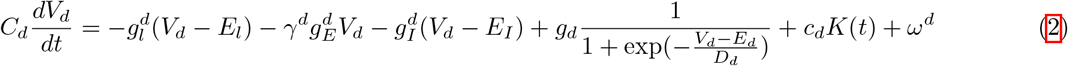

where *C_s/d_* is the somatic/dendritic capacitance, *g_I/E_* is the inhibitory/excitatory conductance (index *d* indicates the dendritic variables), *E_l_* and *E_l_* are the reversal potentials of the leak and the inhibitory synapses, respectively (note that the reversal potential of the excitatory synapses is 0 mV and, therefore, omitted), *τ* is excitability, *ω^d/s^* is the somatic/dendritic adaptation variable. When the soma spikes, the dendrite receives a back-propagating action potential after a delay of 0.5 ms, which is modelled as a 2 ms long current pulse (defined by rectangular kernel *K*(*t*)) with amplitude *c_d_* = 2600pA. With 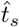 as the time of the last somatic spike, *K*(*t*) is defined as

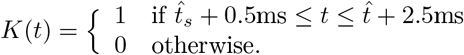

The dendrite has a nonlinear (sigmoidal) term 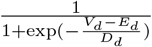 corresponding to the activation of dendritic calcium channels. *E_d_* determines the voltage at which the threshold will be reached and *D_d_* determines the slope of the nonlinear function. The nonlinear dynamics are controlled locally by *g_d_* and are also transmitted to the soma with a coupling factor *g_s_,* such that the soma bursts. The factor λ ensures that the somato-dendritic coupling and adaptation are the same as in the model of Naud and Sprekeler, 2018, where the somatic capacitance was 370 pF (we used *C_s_* =200pF). The somatic adaptation variable is modelled as

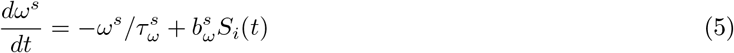

where 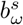, is the strength of spike-triggered adaptation and 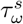, is the recovery time scale. The dendritic adaptation variable is written as

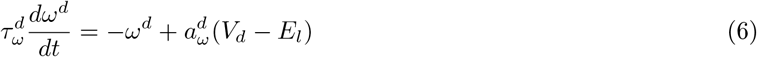

where 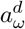 is the strength of subthreshold adaptation and 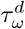 is the recovery time scale.

For the inhibitory population, we used a single-compartment leaky integrate-and-fire neuron model, which membrane potential *V* evolves according to:

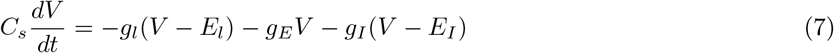

For all neurons, excitatory and inhibitory conductances, *g_E_* and *g_I_* respectively, are increased by the synaptic weight *w_iE_/w_iI_*, depending on their type *i* upon a spike event in a presynaptic excitatory or inhibitory neuron with spike train *S_j_*(*t*), and decay exponentially with time constants *τ_E_* and *τ_i_*, respectively:

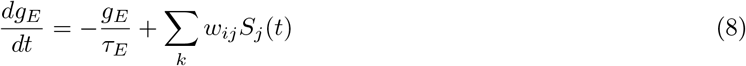

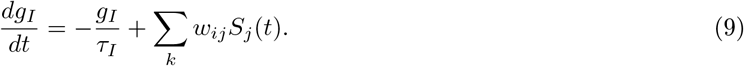

Both excitatory and inhibitory neurons had a refractory period of 8.3 ms (chosen according to the network model from Zenke *et al.*, 2013. Initial membrane potentials for *V_s_* and *V* were sampled from a Gaussian distribution with *μ* = −70mV and *σ* = 10mV to prevent that all neurons spike at the same time at the beginning of a simulation. *V_d_* was set to −70mV initially.

### Plasticity

Synapses from neuron *j* targeting the perisomatic compartment of neuron *i* change their synaptic weight *w_ij_* according to the triplet rule (Pfister and Gerstner, 2006). For clarity we repeat the same equation as in the main text:

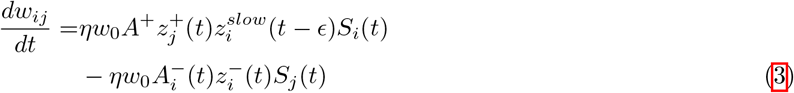

where *w*_0_ is the initial weight, *A*^+/-^ is the amplitude of potentiation/depression (the depression one is time dependent, see below), *S_i/j_* is the post-/presynaptic spike train, 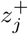 is the presynaptic trace, 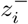 is the postsynaptic trace, 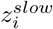 is the postsynaptic trace with a slower time constant. *ϵ* denotes a small fraction of time such that *t* – *ϵ* indicates that the value of the trace is taken before the time point of the action potential *t*. Parameters were chosen as in (Zenke *et al.*, 2013).

Synapses from neuron *j* to neuron *i* targeting the dendritic compartment change their synaptic weight 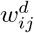 according to the same triplet rule with the back-propagating action potential (bAP) as the postsynaptic spike and an additional Ca-spike-dependent potentiation at the time of a presynaptic spike.

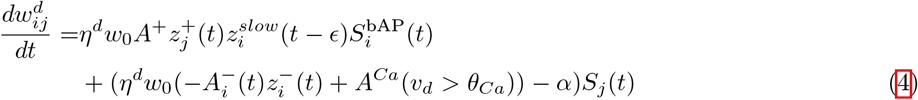

Here, the timing of the back-propagating action potential in the dendrite is used to update the post-synaptic traces 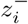 and 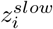 and 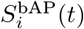 is the postsynaptic train of back-propagating action potentials. A back-propagating action potential is detected if three conditions are met: (1) the dendritic membrane potential *v_d_* exceeds a threshold of −50 mV, and (2) there was a somatic spike within the last 3 ms, and (3) there was no backpropagating action potential within the last 5.8 ms (to account for the refractory period). Synapses are potentiated by a constant amount *A^Ca^* when the presynaptic cell fires and the postsynaptic dendritic membrane potential *v_d_* exceeds a threshold *θ_Ca_* of −40mV. The term *v_d_* > *θ_Ca_* takes a value of 1 when the threshold is crossed and is 0 otherwise. Synapses are depressed by a constant amount *α* for each presynaptic spike (transmitter-induced plasticity).

The pre- and postsynaptic traces are defined as:

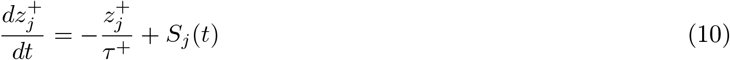

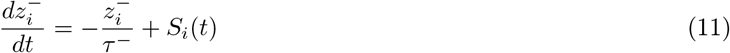

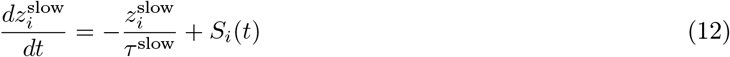

where *τ*^+^, *τ*^-^, and τ^slow^ are the time constants with which the traces decay. Both perisomatic and dendritic excitatory synapses are limited by a maximum synaptic weight *w*_max_ = 10 nS.

### Homeostatic Plasticity

The depression amplitude 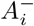 for all synapses onto neuron *i* is a function of a moving average of neuron *i*’s activity 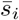:

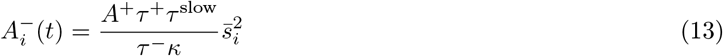

where *κ* is the target firing rate, *A*^+^, *τ*^+^, *τ*^-^ and *τ*^slow^ are variables from the triplet STDP rule and 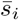 is the low-pass filtered spike train:

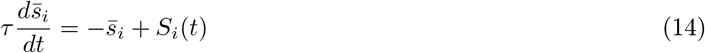

with *τ* defining the time constant of the homeostatic plasticity.

### Synaptic Scaling

In Suppl. Fig. S5, instead of a sliding depression amplitude (Bienenstock *et al.*, 1982), we used synaptic scaling as a homeostatic mechanism, implemented by the term 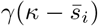 in the following weight update equations. As before 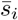 is the moving average of neuron *i*’s activity, *κ* is the target firing rate. For perisomatic synapses, weights change according to:

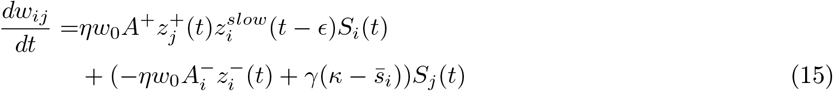

where

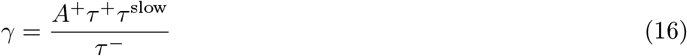

Equivalently, for the dendritic synapses, weights change according to:

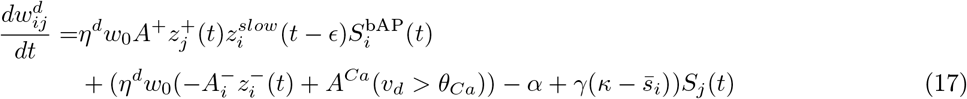

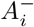 was fixed to 2.7e-3.

### Explosion factor

We quantify the stability with the explosion factor EF. We calculate it as follows:

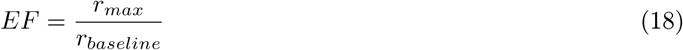

where *r_max_* is the maximum population firing rate within the duration of the simulation and *r_baseline_* is the population firing rate averaged over the first 50 ms of the simulation. Therefore, an explosion factor close to 1 indicates that the network activity is stable. The distribution of explosion factors was bimodal with a sharp peak close to 1 and a broader distribution of larger EFs (Fig. 2h). We defined a threshold separating those two modes, which defines whether the network is stable or explodes:

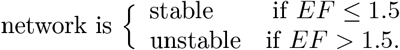

### Critical time constant *τ_crit_*

For each value of the gate, we calculated the maximum *τ* for which the network was stable. It additionally had to be smaller than the minimum *τ* for which the network was unstable.

### Baseline dendritic weight change

For each gating value, we calculated the sum of all weight changes in the dendrite in a 200 s simulation with a *T* of 5 ms.

### Statistical analyses

To test for significance in Fig. 3 and Fig. 4, we used the two-sample student’s t-test.

### Simulation

For Fig. 1, we simulated the network for 10s without plasticity. Simulations to calculate the explosion factor (for Figs. 2-4) were run for 200 seconds. We simulated an initial warm-up phase for 3 *τ* seconds without plasticity to calculate the average population firing rate for the balanced network. We used the average population firing rate of the last 2 seconds of the warm-up phase to set the target firing rate *κ* in our model. We then switch on plasticity. All simulations were run at a timestep of 0.1 ms. For the plots in Figs. 1-3, each condition was simulated with 10 different seeds.

### Memory network (Fig. 5)

We used the described network and added plasticity on inhibitory to excitatory synapses and a competition mechanism for postsynaptic weights.

#### Inhibitory Plasticity

Synapses from inhibitory neuron *j* to excitatory neuron *i* change their weight 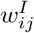 according to Vogels *et al.*, 2011

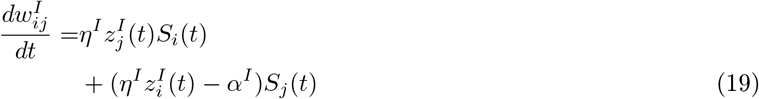

where *η^I^* is the inhibitory learning rate, 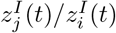 is the pre/post-synaptic trace, *S_j_*(*t*)/*S_i_*(*t*) is the pre/post-synaptic spike train, and *α^I^* = 2*κT*_*iSTDP*_ determines the amount of transmitter-induced depression Vogels *et al.*, 2013. Plastic inhibitory weights are limited by a maximum weight 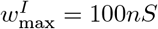. The pre/post-synaptic traces 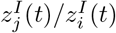 are written as

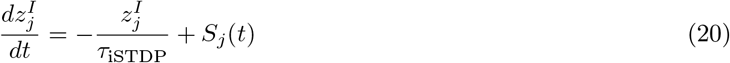

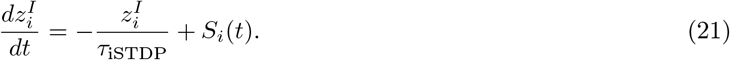

For Suppl. Fig. S4, we explored a different form of inhibitory plasticity with an asymmetric learning rule:

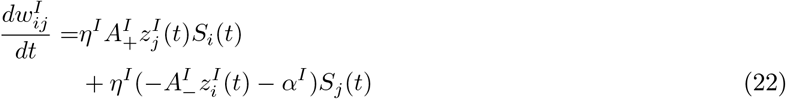

where 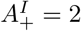 and 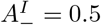.

#### Heterosynaptic depression

If the sum of postsynaptic weights in the perisomatic compartment or the dendrite exceeds a maximum 1.5*pN_E_w*_0_ (hard bound), all perisomatic/dendritic synaptic weights are scaled down equally by the average synaptic weight change of the postsynaptic compartment in the current time step, that is the total perisomatic/dendritic weight change divided by the number of incoming perisomatic/dendritic synapses.

#### Stimulation protocol

We first simulated the non-plastic network for 5 s to calculate the steady state population firing rate. We then introduced plasticity with a target firing rate *κ* of the measured population firing rate and simulated the network with plasticity for another 5 s. Then, we activated pattern P1 for 3 s, which was realised by an external input of 100 Poisson spike trains with a firing rate of 20 Hz to neurons with indices 400 to 499 (ensemble E1). Afterwards, we changed the gating variable under investigation (except in Fig. 5 a, where we did not apply any gating). After a stimulation pause of 7 s, we activated pattern P2 for 5 s, which was realised by an external input of 100 Poisson spike trains with a firing rate of 30 Hz to neurons with indices 450 to 549 (ensemble E2). We continued the simulation for further 10 s without stimulation.

#### Single-compartment network

For Suppl. Fig. S7, we simulated a network of the same size, where excitatory cells were single-compartment neurons. The somatic membrane equation was the same as for the two-compartment neurons with *g_s_* =0 (no coupling to the dendrite). All parameters were the same as in the original network, with two exceptions, as we placed all synapses on the perisomatic compartment. First, the connection probability from excitatory to excitatory cells was increased to *p_EE_* = 0.19 to account for the additional synapses that were previously placed on the dendrite. Second, in addition to the inhibitory synapses the soma already contained in the original simulation, it contained inhibitory synapses with the connection strength of inhibitory synapses on the dendrite in the original model 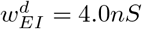 with a probability of 0.1.

#### Parameters

**Table 1:** Parameters of the network.

| Parameter | Value |
| --- | --- |
| $N_E$ | 1000 |
| $N_I$ | 250 |
| $N_{\text{Poisson}}$ | 1000 |
| $\lambda_{\text{Poisson}}$ | 2 Hz |
| $p$ | 0.1 |
| $w_{EP}$ | 1.6 nS |
| $w_{IP}$ | 1.6 nS |
| $w_{EE}$ | 1.8 nS |
| $w_{EE}^d$ | 1.8 nS |
| $w_{IE}$ | 4.0 nS |
| $w_{II}$ | 6.0 nS |
| $w_{EI}$ | 8.0 nS |
| $w_{EI}^d$ | 4.0 nS |

**Table 2:** Parameters of the neuron model.

| Parameter | Value |
| --- | --- |
| $g_l$ | 10.0 nS |
| $g_l^d$ | $\frac{170}{7}$ nS |
| $E_l(\text{leak})$ | -70 mV |
| $E_I(\text{Inhibitory})$ | -80 mV |
| $v_{\theta}$ | -50.0 mV |
| $v_{\theta}^I$ | -50 mV |
| $\tau_m$ | 20 ms |
| $\tau_m^I$ | 10 ms |
| $C_s$ | 200 pF |
| $C_d$ | 170 pF |
| $c_d$ | 2600 pA |
| $a_{\omega}^d$ | -13 nS |
| $b_{\omega}^s$ | -200 pA |
| $\tau_{\omega}^s$ | 100 ms |
| $\tau_{\omega}^d$ | 30 ms |
| $\tau^s$ | 16 ms |
| $\tau^d$ | 7 ms |
| $g_s$ | 1300 pA |
| $g_d$ | 1200 pA |
| $E_d$ | -38 mV |
| $D_d$ | 6 mV |
| $\lambda$ | 0.54 |

**Table 3:** Parameters of the plasticity.

| Parameter | Value |
| --- | --- |
| $A^+$ | 6.5e-3 |
| $A^{Ca}$ | 7.2e-2 |
| $\alpha$ | 1e-4 |
| $\theta_{\text{bAP}}$ | -50 mV |
| $\theta_{Ca}$ | -40 mV |
| $\tau^+$ | 16.8 ms |
| $\tau^-$ | 33.7 ms |
| $\tau^{\text{slow}}$ | 114 ms |
| $w_{\text{max}}$ | 10 nS |
| $\tau_{\text{iSTDP}}$ | 20 ms |
| $w_{\text{max}}^I$ | 100 nS |

## Data and material availability

The simulation code will be made available on GitHub at https://github.com/k47h4/gating_plasticity upon acceptance of the manuscript.

## List of supplementary material

- S1: Dendritic weight changes as a function of *τ_crit_* in the network simulations shown in Fig. 2e
- S2: Effect of dendrite-to-soma coupling and the dendritic nonlinearity. Supplementary Figure to section 3.
- S3: Additional plots for the memory network. Supplementary Figure to section 5.
- S4: Protection of memories by gating plasticity in a network with asymmetric inhibitory plasticity. Supplementary Figure to section 5.
- S5: The network with synaptic scaling as a homeostatic mechanism.
- S6: Dependence of the critical homeostatic time constant on parameters.
- S7: Comparison of gating plasticity in the dendrites in a network with two-compartment neurons to gating plasticity in the perisomatic compartment in a network with single-compartment neurons.

## Supplementary Figures

**S 1.**
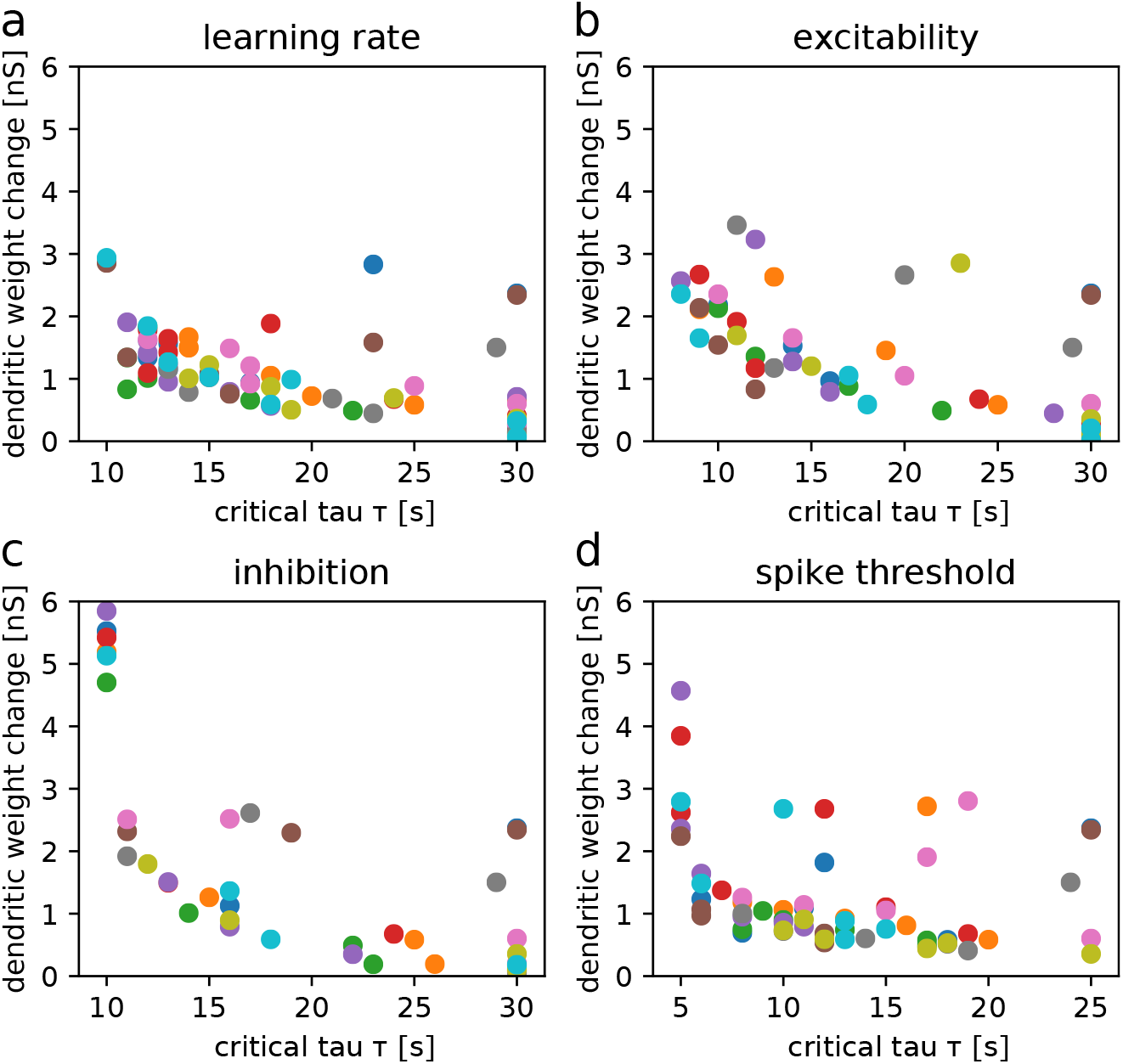
Dendritic weight changes as a function of *r_crit_* in the network simulations shown in Fig. 2e where learning rate (a), excitability (b), inhibition (c), and spike threshold (d) were varied. The different colours represent the 10 different seeds used for the simulations.

**S 2.**
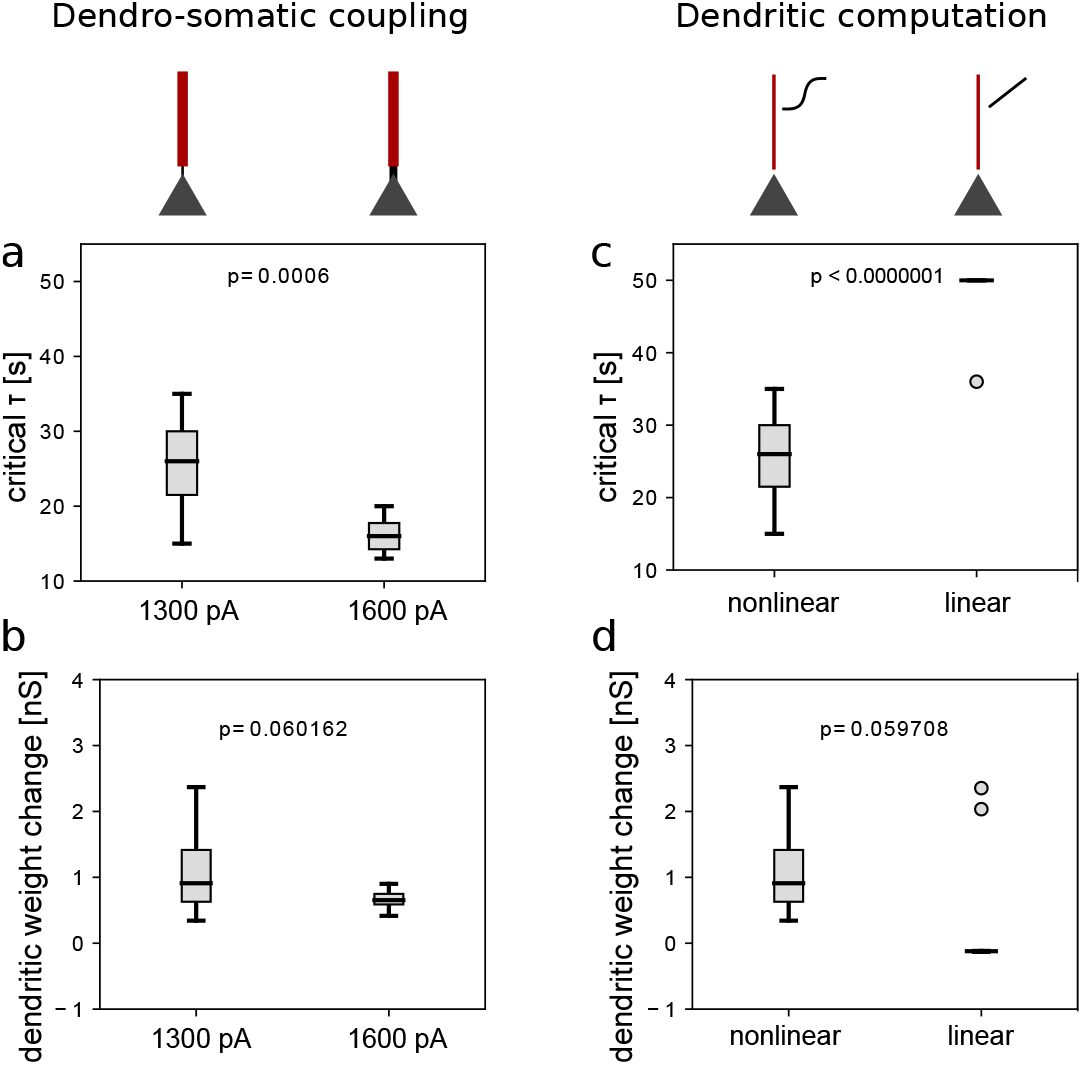
Effect of dendrite-to-soma coupling and the dendritic nonlinearity. a,b: Effect of the dendrite-to-soma coupling, determined by *g_s_,* i.e. how much the dendritic nonlinearity affects the soma. a: Distribution of critical homeostatic time constants for a dendro-somatic coupling of 1300 pA (default) versus 1600pA. b: Distribution of dendritic weight changes for a dendro-somatic coupling of 1300 pA (default) versus 1600pA. c-d: Effect of the dendritic nonlinearity. c: Distribution of critical homeostatic time constants for a network with nonlinear dendrites and a network with linear dendrites. d: Distribution of dendritic weight changes for a network with nonlinear dendrites and a network with linear dendrites. The rectangles represent the interquartile range (IQR) between first and third quartiles. The thick horizontal lines represent the medians. The whiskers indicate the lowest and highest values within 1.5xIQR from the first and third quartiles, respectively. The circles denote outliers. p-values were obtained by using the two-sample student’s t-test.

**S 3.**
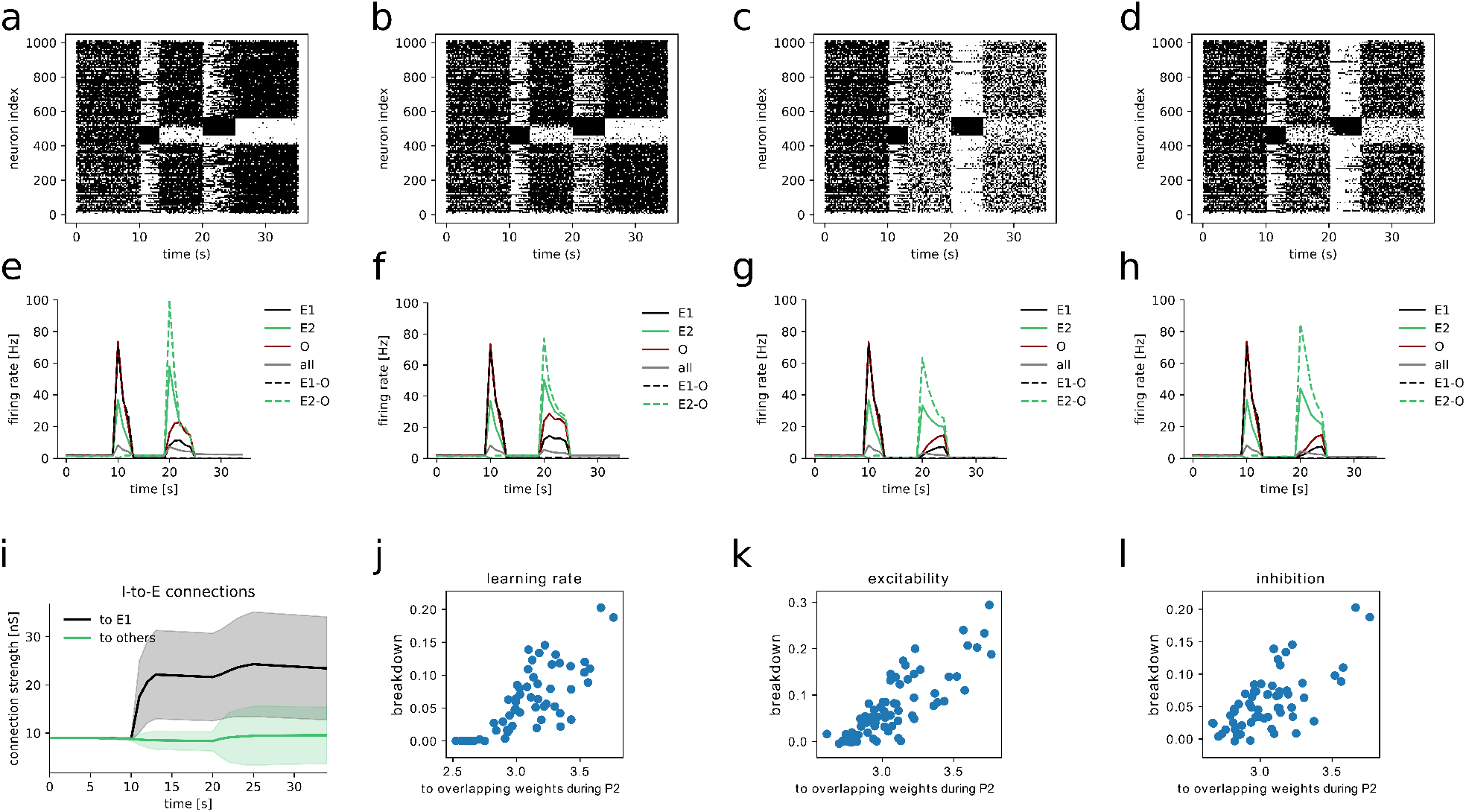
Additional plots for the memory network. a-d: Raster plots for the simulation without gating (a), with gated learning rate (b), with gated excitability (c) and with gated inhibition (d) as in Fig. 5. e-h: Firing rates over time of different subpopulations: E1: memory ensemble E1 (*x* ∈ *E*1), E2: memory ensemble E2 (*x* ∈ *E*1), O: overlapping ensemble O (*x* ∈ (*E*1 ∩ *E*2), all: all excitatory neurons in the network, E1-O: ensemble E1 excluding the overlapping ensemble O (*x* ∈ (*E*1 \ *O*)), E2-O: ensemble 2 excluding the overlapping ensemble (*x* ∈ (*E*2 \ *O*)). i: Connection strengths over time from the inhibitory population to ensemble E1 (black) and from the inhibitory population to all other cells in the ungated network. j-l: Memory breakdown as a function of the maximum during pattern P2 of the mean strength of synaptic connections from all excitatory cells to the overlapping ensemble O for a network with gated learning rate (j), gated excitability (k) and gated inhibition (l).

**S 4.**
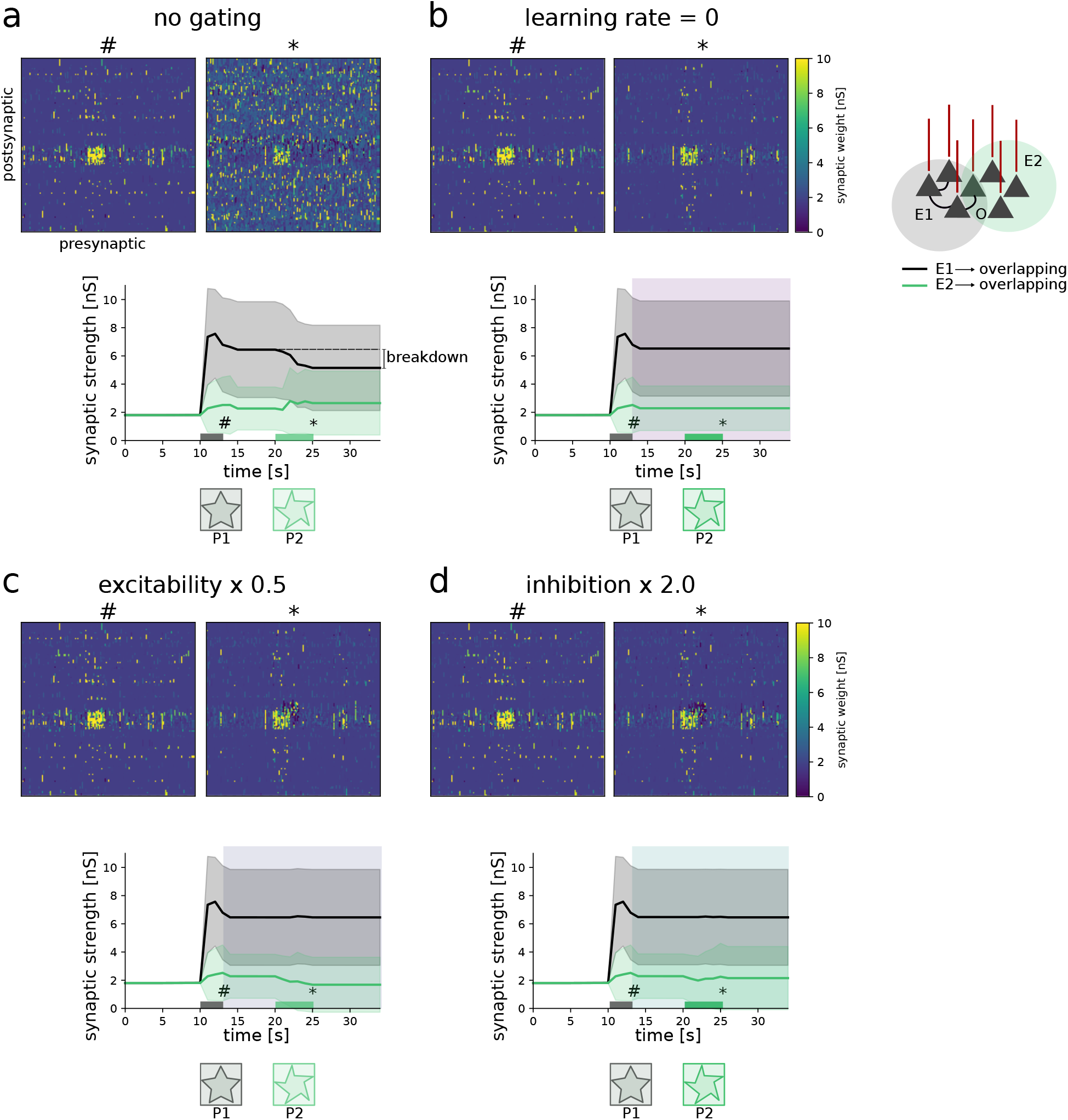
Protection of memories by gating plasticity in a network with asymmetric inhibitory plasticity. a-d: After 10 seconds, we show pattern 1 (P1, grey) to the network for 3 seconds. After a gap of 7 seconds, we show pattern 2 (P2, green) to the network for 5 seconds. Top panels: excitatory weight matrix at two time points of the simulation. Bottom panel: mean synaptic weight from ensemble 1 (E1) to the overlapping region (grey) and from ensemble 2 (E2) to the overlapping region over time as an indication of the strength of the memory of P1 and P2, respectively. a: no gating. b: after P1 is learned, the learning rate is set to 0 (denoted by purple background). c: after P1 is learned, excitability is reduced by 50% (denoted by blue background). d: after P1 is learned, inhibition is doubled (denoted by green background).

**S 5.**
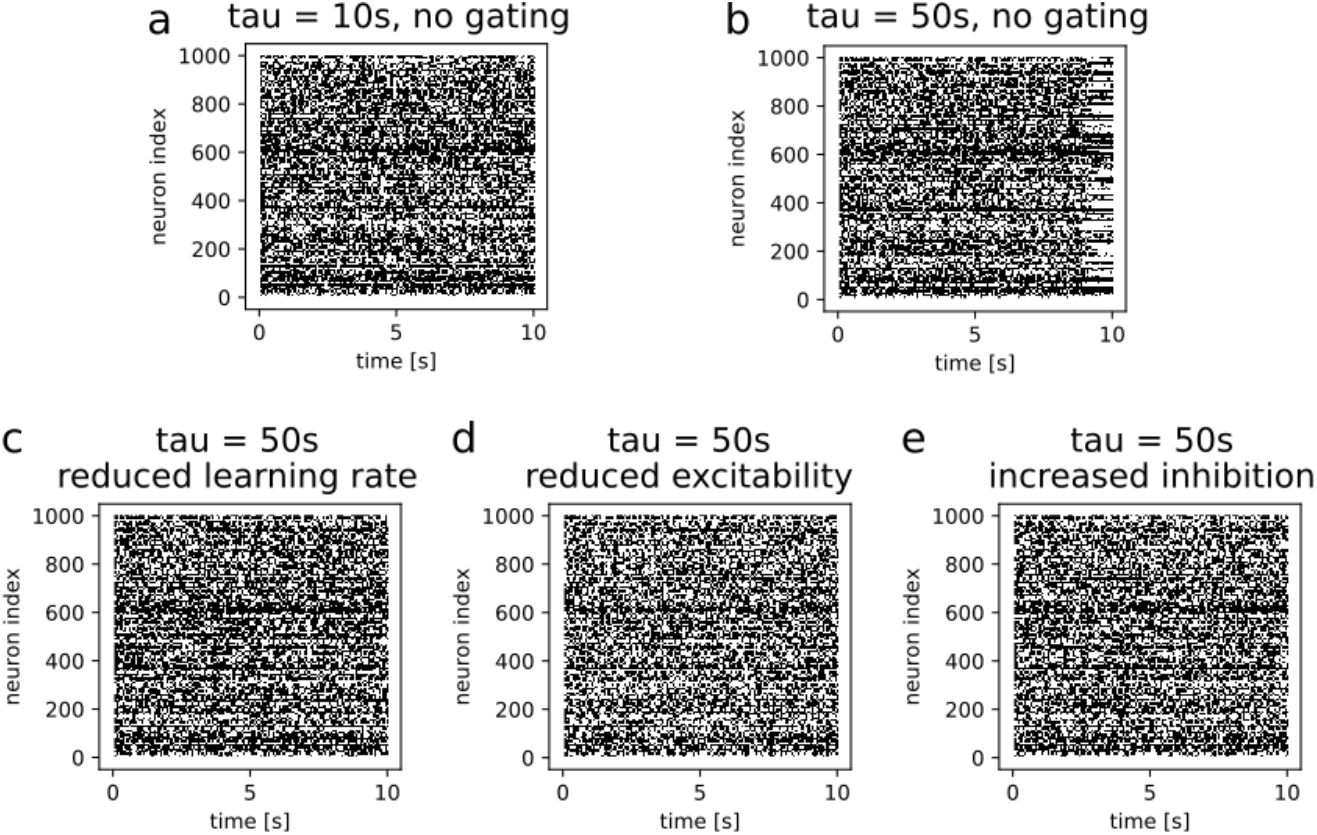
The network with synaptic scaling as a homeostatic mechanism. Raster plots of a network with synaptic scaling as a homeostatic mechanism. a: no gate applied, synaptic scaling with a homeostatic time constant of 10s. b-e: synaptic scaling with a homeostatic time constant of 50s. b: no gate applied. c: reduced learning rate (reduced by 40%). d: reduced excitability (reduced by 10%). d: increased inhibition (increased by 10%).

**S 6.**
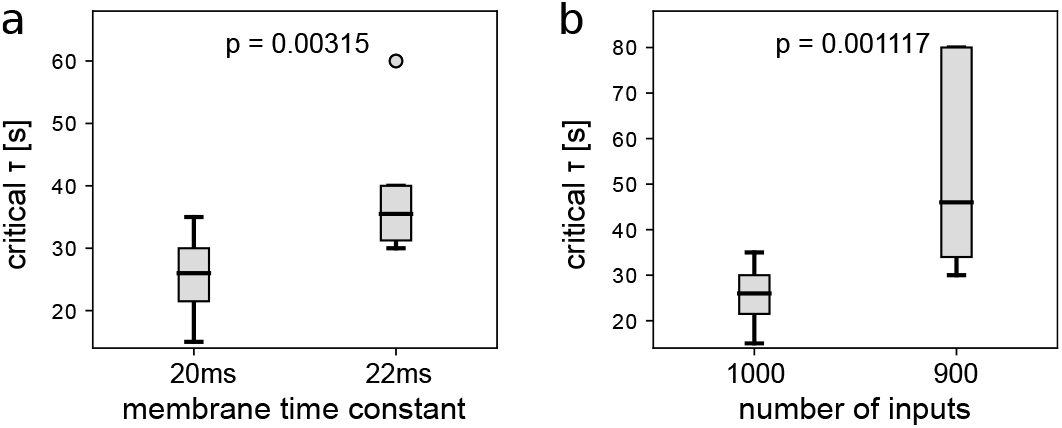
Dependence of the critical homeostatic time constant on parameters. a: The network was simulated with two different membrane time constants. For each condition, the network was simulated 10 times with different seeds. The distribution of critical homeostatic time constants is plotted for the network with a membrane time constant of 20ms, and for the network with a membrane time constant of 22ms. b: The network was simulated with different numbers of excitatory Poisson inputs. Both the excitatory and the inhibitory cells receive these inputs. The distribution of critical homeostatic time constants is shown for the network with 1000 Poisson inputs, and for the network with 900 Poisson inputs. The rectangles represent the interquartile range (IQR) between first and third quartiles. The thick horizontal lines represent the medians. The whiskers indicate the lowest and highest values within 1.5xIQR from the first and third quartiles, respectively. The circles denote outliers. p-values were obtained with the two-sample student’s t-test.

**S 7.**
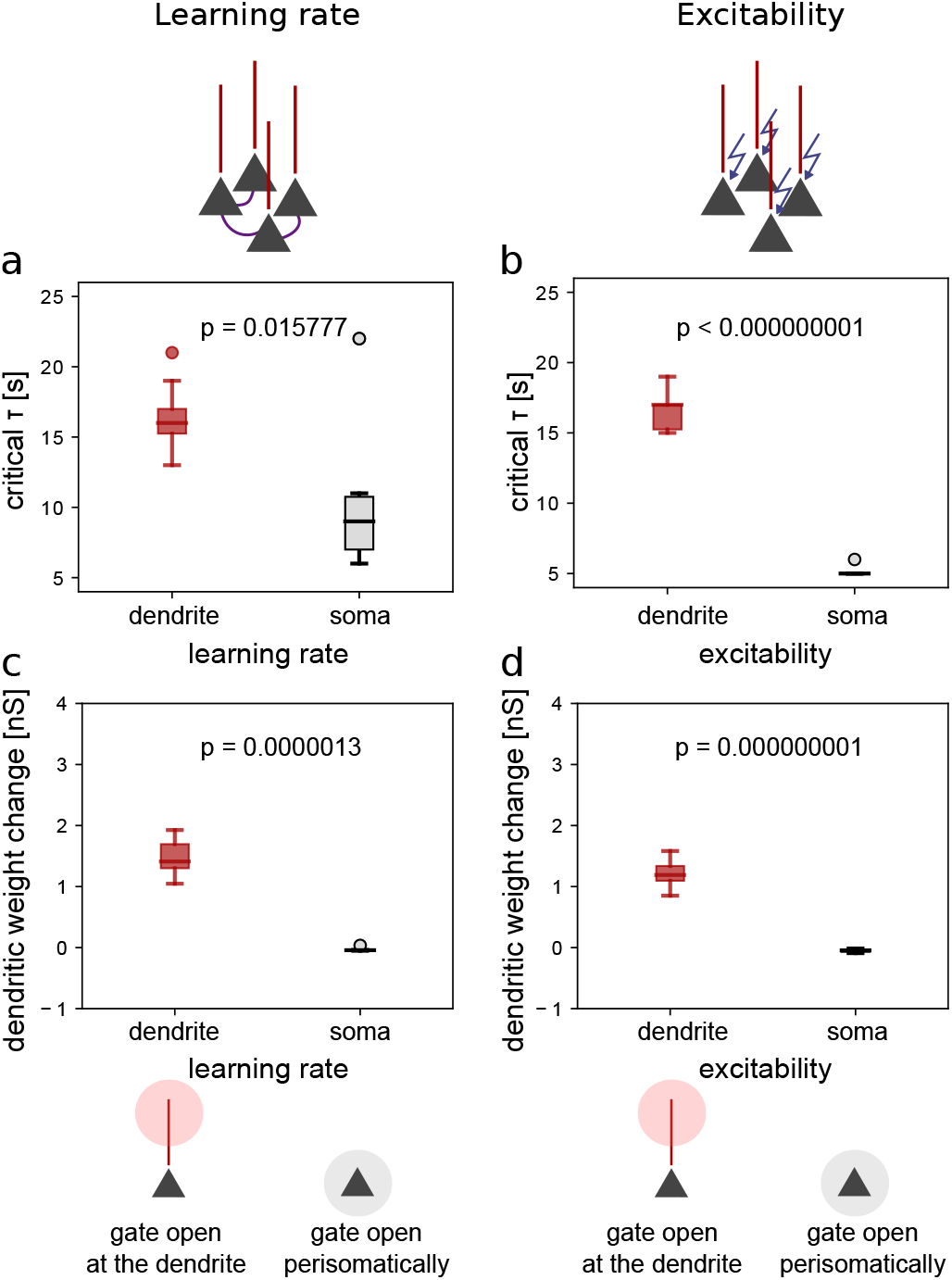
Comparison of gating plasticity in the dendrites in a network with two-compartment neurons to gating plasticity in the perisomatic compartment in a network with single-compartment neurons. a-b: Distribution of critical homeostatic time constants for gating in the dendritic (red) and in the perisomatic (black) synapses for (a) a two-fold increase in the learning rate, and (b) a 15% increase in excitability. c-d: Distribution of dendritic weight changes for gating in the dendritic (red) and in the perisomatic (black) synapses for (c) a two-fold increase in the learning rate, and (d) a 15% increase in excitability. The rectangles represent the interquartile range (IQR) between first and third quartiles. The thick horizontal lines represent the medians. The whiskers indicate the lowest and highest values within 1.5xIQR from the first and third quartiles, respectively. The circles denote outliers. All p-values were obtained by using the two-sample student’s t-test.

